# Visual features underlying neural selectivity for natural stimuli

**DOI:** 10.64898/2025.12.08.692716

**Authors:** Jan Kevin Schluesener, Malav Shah, Florian Sandhaeger, Markus Siegel

## Abstract

Neural activity is selective for the identity and semantic category of natural visual stimuli. In human neuroimaging, this selectivity can be investigated using multivariate pattern analysis. The resulting neural information is thought to reflect neural processes underlying object recognition. However, due to the complexity of natural stimuli, it remains unclear to what extent different visual features drive this information. To address this, we presented natural stimuli from different categories as well as several degraded versions, in which specific visual features were selectively manipulated, to human participants while recording magnetoencephalography (MEG). We applied multivariate pattern analysis and decomposed the resulting high-dimensional neural representational geometry into its distinct contributions from retinotopy, total spatial frequency and orientation energy, global shape, local shape, and textural correlation structure. These features contributed differentially to the neural information, with retinotopy and global shape being the strongest features underlying image and category information, respectively. Our results demonstrate how multiple statistical features of visual stimuli jointly and dynamically account for neural image and category selectivity, providing a basis for a better understanding of human object recognition.

## Introduction

When presented with natural visual stimuli, the human brain extracts low-dimensional semantic information from retinal inputs. This process relies on the distribution of low-level visual features, such as color, orientation, or spatial frequency, which, through a cascade of processing steps, are reformatted into more complex, semantic features. As this process unfolds, neural activity becomes selective for both the identity and category of stimuli. In human neuroimaging, this selectivity can be captured by multivariate decoding methods [1,2], and the emergence of neural stimulus or category information is usually interpreted as a signature of the processing, categorization, and recognition of the stimulus, which ultimately allows us to select an appropriate behavioral response.

The mechanisms of how we can recognize and categorize visual inputs are not fully understood. Artificial neural network models, which are able to perform these tasks at levels comparable to humans, have been proposed as a promising model of the underlying mechanisms [3]. However, while advances in neural network modelling have enabled impressive accuracies in object recognition tasks, these artificial models can behave subtly different from humans [4–7]. Furthermore, they are not sufficiently constrained to uncover the biologically implemented processes underlying human object recognition [8].

An important element that is missing to gain this understanding is a full characterization of the dynamic contributions of visual features driving neural selectivity. For simple visual stimuli, it is possible to quantify the relative contribution of multiple low-level features to overall neural activity and stimulus selectivity [9–11]. However, for more complex natural stimuli it is challenging to simultaneously control and quantify the contribution of all low-level visual features.

An alternative approach consists in the use of large, high-dimensional batteries of stimuli and the subsequent comparison of neural representational spaces with various feature-based representational models [12–16]. While this can successfully uncover a relationship between neural signals and many visual or semantic features, complex inter-dependencies within the feature space make it difficult to conclusively quantify the unique contribution of individual features. Using stricter experimental control, several studies have shown how the selectivity for categories of natural stimuli depends on individual visual features, such as shape [17–19], texture [19], or spatial frequency [20]. Here, we aimed to extend this approach to comprehensively characterize the contribution of the constitutive statistical features of natural stimuli to the selectivity for both individual images and semantic categories.

To this end, we devised a stimulus set allowing us to quantify the effect of five different statistical features on image and category selectivity in human magnetoencephalography (MEG) recordings. The stimulus set consisted of natural visual stimuli of objects belonging to different categories and multiple degraded versions of each stimulus. Using a multivariate pattern analysis approach, we decomposed the representational geometry of this high-dimensional stimulus space. Specifically, the employed image degradations enabled us to isolate the contributions of retinotopy, total spatial frequency and orientation energy (the power spectrum), global shape [21], local shape, and textural correlation structure [22]. The construction of the stimulus space implicated that these five features fully described each visual stimulus. We found that the neural information about individual images was initially dominated by retinotopic differences, while contributions of global and local shape as well as orientation and spatial frequency energy gradually developed later. In contrast, the neural information about semantic categories was dominated by global shape, with only minor contributions of the other features. Overall, we show how multiple statistical features of visual stimuli jointly and dynamically account for neural image and category selectivity.

## Results

### MEG recordings during presentation of natural and modified images

We recorded MEG in 18 human participants that viewed a stream of original and modified natural images while they performed a visual target detection task, that ensured attentive processing of the visual stimuli (Fig. 1a). On each trial, a random sequence of stimuli of unpredictable length was presented. Participants had to maintain fixation and continually press a button until a small visual go cue appeared at an unpredictable time and location, instructing them to release the button.

**Fig 1.**
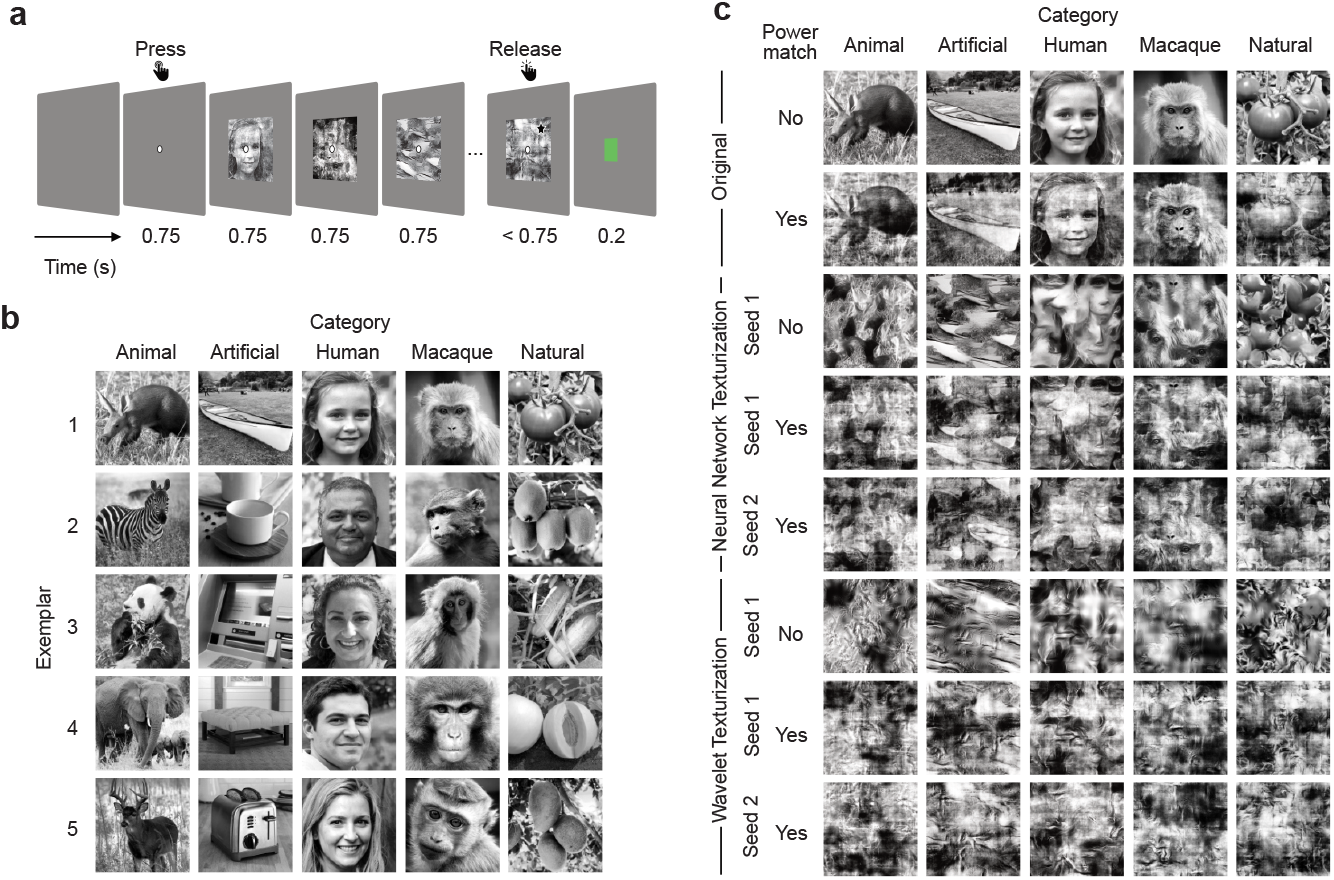
Behavioral task and stimuli. (**a**) Participants initiated each trial by button press. A central fixation spot appeared, which subjects fixated on throughout the trial. Following a blank baseline (0.75 s) a sequence of images (0.75 s each) was presented. After an unpredictable number of stimuli, a go cue (a black star) appeared at an unpredictable location on the last stimulus. Subjects had to release the button, within the duration (0.75 s) of the last stimulus. Visual feedback (a green or red square) was provided after each trial. (**b**) 25 natural grayscale images from 5 different categories (artificial objects, natural objects, animals, macaques, human faces) (**c**) Original images (top row) and their 7 modifications for exemplar image 1 of each category. Images were modified using three approaches: spectral power matching (PM), wavelet texturization (WT) and neural network texturization (NNT). For power matched stimuli, NNT and WT texturizations were performed twice with different seeds. Human face images were artificially generated using a generative adversarial neural network model [55] (https://thispersondoesnotexist.com/).

Each subject viewed a set of 200 stimuli, consisting of 25 natural grayscale images from 5 different categories (artificial objects, natural objects, animals, macaques, human faces) (Fig. 1b) as well as 7 different modifications of each stimulus (Fig. 1c). Modified stimuli were generated using a combination of three strategies: First, to investigate the effects of total spatial frequency and orientation energy, we created power-matched stimuli, for which the 2D Fourier spectra of all 25 images were equalized (Fig. 1c; power match). Second, to disentangle effects of global shape, local shape, and texture as defined by the textural correlation structure [22], we employed two different texturization algorithms. The first one was based on a neural network approach disrupting global shape but preserving texture and local shape elements [21] (Fig. 1c; neural network texturization). The second was a parametric wavelet-based texture model disrupting both global and local shape, and preserving only texture [22] (Fig. 1c; wavelet texturization). Third, to isolate the effect of retinotopic shifts of image features, we applied each texturization algorithm twice with different starting conditions, resulting in images with the same texture and/or local shape features, but different spatial arrangements (Fig. 1c; seed 1 vs seed 2). Notably, a common definition of texture would include both the textural correlation structure and the power spectrum [22]. Combining spectral power-matching with wavelet texturization allowed us to dissociate these two features. Furthermore, to discount potential confounds of luminance contrast the luminance histogram of all original and modified images was matched to the average histogram of all original images.

Through this combination of approaches, we arrived at the final set of modifications: original images (Original), original power matched images (Original PM), neural-network texturized images (NNT), two versions of neural-network texturized and power matched images (NNT 1 PM and NNT 2 PM), wavelet-texturized images (WT), and two versions of wavelet-texturized and power-matched images (WT 1 PM and WT 2 PM). To rule out spurious results due to idiosyncrasies of the specific images chosen, we created two different sets of 25 original stimuli and used set 1 and set 2 for half of the subjects each.

### Extracting neural imageand category information

We used multivariate pattern analysis to extract the information contained in MEG signals about both the identity and the semantic category of the presented images. First, we computed a time-resolved, pairwise dissimilarity matrix between all stimuli using the cross-validated Mahalanobis distance [23] Fig. 2a). We calculated image information as the mean pairwise distance between image pairs from the same category and modification (Fig. 2a left). To extract category information, for each modification, we used the mean pairwise distances between images from different categories (Fig. 2a middle) and subtracted image information from this value. Thus, neural image information quantified the discriminability of neural responses to different natural stimuli within a semantic category; whereas neural category information quantified the discriminability of neural responses to different semantic categories, independent of the specific images displayed.

**Fig 2.**
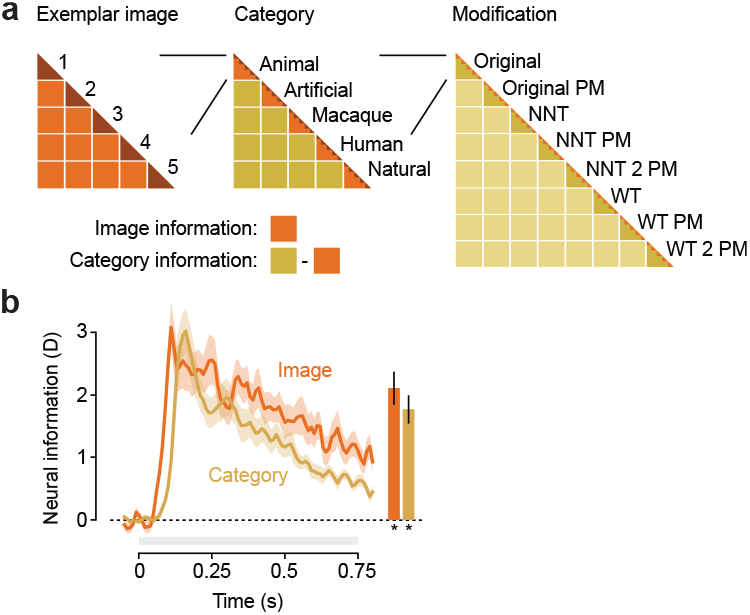
Neural dissimilarity matrices and information. (**a**) The full dissimilarity for all pairs of stimuli can be partitioned into dissimilarities between images of the same category and modification (left), between images of different categories of the same modification (middle), and between images of different modifications (right). (**b**) Neural image and category information for original images. Image information is the crossvalidated Mahalanobis distance (D) between images of the same category. Neural category information is the cross-validated Mahalanobis distance (D) between images of different categories minus the distance between images of the same category. The bars on the right show the average information 0.15 to 0.5 s post stimulus onset. Asterisks below bars indicate significant information (p < 0.05). All error bars indicate the SEM across subjects.

Applying this method to the original natural stimuli yielded robust neural information about both, image identity and category (Fig. 2b). Both image and category information emerged rapidly after stimulus onset and slowly decreased during stimulus presentation. Image information was of similar magnitude (two-sided t-test, p = 0.0879, 0.15 to 0.5 s) but increased earlier (time at half-maximum: 83 ± 3 ms vs. 118 ± 4 ms, p = 6*10^−8^) than category information.

### Equating spectral power reduces neural information but preserves representational structure

How does the removal of differences in spectral power, i.e. of the total spatial frequency and orientation energy, affect neural responses? As MEG signals reflect the average activity of large neural populations, their selectivity may be dominated by these summary statistics. To investigate this, we compared neural image and category information between original and original power-matched stimuli. Both types of information were reduced by power matching (Fig. 3a, b; image: p = 0.001; category: p = 0.0003), indicating that neural representations became more similar by removing spectral differences. Still, both types of information were significant after power matching (Fig. 3a, b; image: p = 8*10^−7^; category: p = 3*10^−8^).

**Fig 3.**
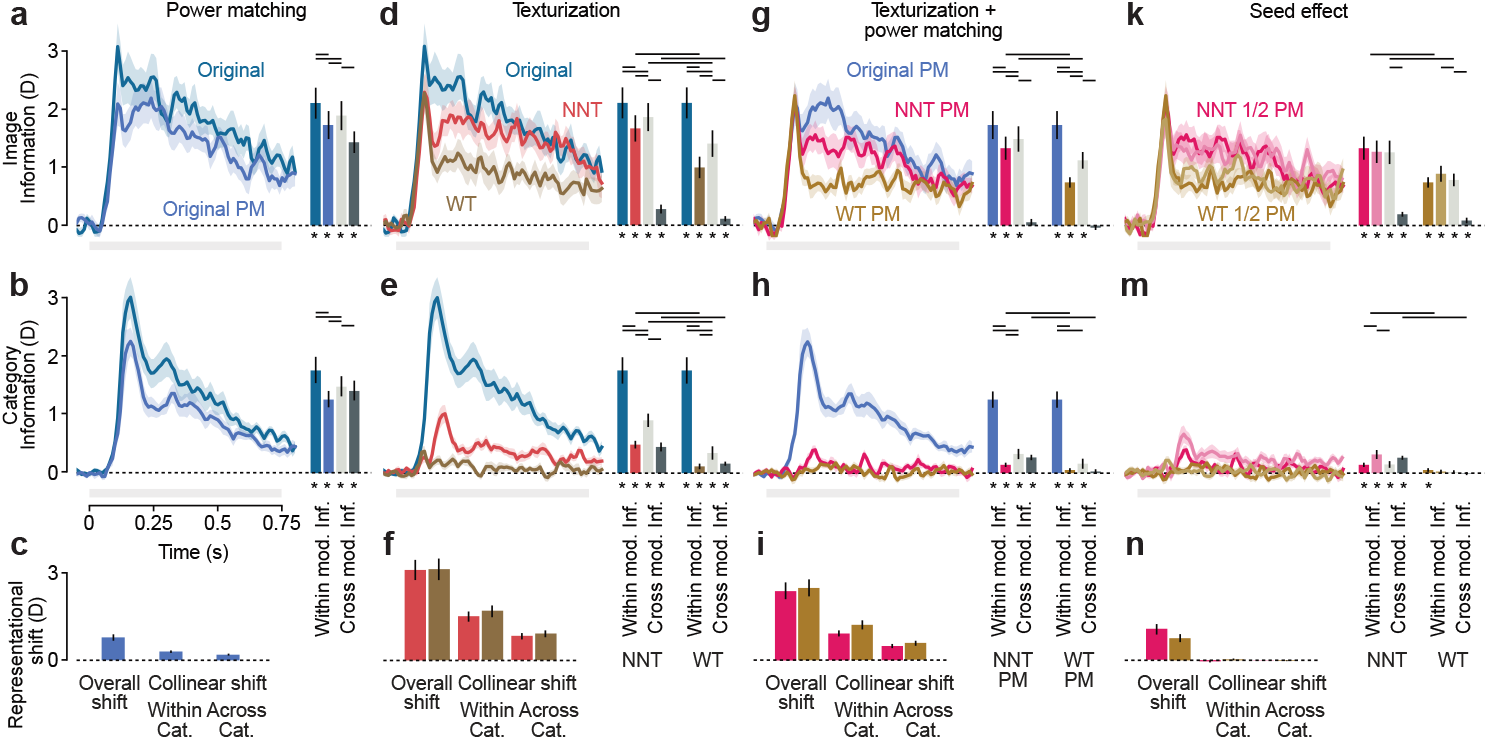
The effect of image modifications on neural image and category information. (**a**) Neural image information for original and power matched original images. The colored bars on the right show the average information 0.15 to 0.5 s post stimulus onset. The dark gray bar shows the crossdecoding of image information between original and power matched original stimuli. The light gray shows the cross-modification information expected for representations of original and power-matched stimuli, that only differ in strength but not in the underlying multivariate patterns. (**b**) As in (a) for category information. (**c**) Left: cross-validated Mahalanobis distance (D) between original and power-matched versions of each image. Middle: Collinear cross-validated Mahalanobis distance (D) between different original and power-matched images of the same category. Right: Collinear cross-validated Mahalanobis distance (D) between original and power-matched images of different category. (**d, e, f**) as (a, b, c) comparing neuronal network texturized, wavelet texturized and original stimuli. (**g, h, i**) as (d, e, f) comparing power matched stimuli. (**k, m, n**) as (a, b, c) comparing two different version of power matched neuronal network texturized and power matched wavelet texturized stimuli. Asterisks below bars indicate significant information (p < 0.05). Horizontal lines between bars indicate significant differences information (p < 0.05). All error bars indicate the SEM across subjects.

While the total spectral power was identical for all power-matched images, both local low-level features such as orientation or contrast and semantics were subjectively largely preserved. Thus, we next asked whether there were shared neural representations of original and power-matched images or categories. We used a cross-decoding approach: we computed cross-validated Mahalanobis distances using a pair of original stimuli as training data and the same pair of power-matched stimuli as test data, and vice versa (Fig. 3a, b dark gray bars). In addition, we computed a measure of the maximally expected cross-information under the assumption of identical representations of original and power-matched stimuli, that only differ in strength but not in the underlying multivariate pattern (Fig. 3a, b light gray bars) [24]. This allowed us to test whether the generalization between both modifications deviated from this null hypothesis of identity. We found that both image and category information were largely shared between original and power-matched stimuli (Fig. 3a, b dark gray bars; cross-information > 0: image: p = 3*10^−7^, category: p = 1*10^−7^). Nonetheless, in both cases, generalization was weaker than expected for identical representations (Fig. 3a, b light vs. dark gray bars; cross-information < expected: image: p = 6*10^−5^, category: p = 0.007). While significant, these differences between empirical and expected cross-information were small. Thus, neural representations of images and semantic categories were mostly, but not perfectly preserved by power-matching. Furthermore, generalization was stronger for category information than for image information (empirical cross-information divided by cross-information expected for identical representations: image = 0.754, category = 0.941, p = 0.001). Thus, category information was more invariant to spectral changes than image information.

Original and power-matched images are subjectively clearly distinct. In line with this, both the strong, but imperfect generalization, and the reduction in overall information show a change in neural responses due to power-matching. To quantify the overall magnitude of this change, we computed the neural distance between original and power-matched versions of each image. We found that this representational shift was significant (Fig. 3c, left p = 8*10^−7^), indicating separable neural representations of an original image and its power-matched version.

To further understand the quality of this shift within and across categories, we computed its collinearity across all image pairs, again using a cross-decoding approach (Fig. 3c middle and right; see Methods for further details). This revealed a significant collinearity both within (Fig. 3c, middle p = 1*10^−6^) and across categories (Fig. 3c, right p = 6*10^−6^). Thus, power-matching did not have an independent effect on the neural representation of each image, but led to a structured shift in representational space. The collinearity of this shift was more pronounced within categories than across (p = 0.003), suggesting that stimuli within the same category partially share their overall spectral power and consequentially undergo a similarly directed shift towards a common mean due to power-matching.

### Disrupting shapes strongly attenuates category information

Next, we assessed the effect of local and global shape on neural information. We compared neural responses for original natural stimuli and texturized stimuli that had all shape features removed (wavelet texturization; WT) and those that had all global shape removed but retained some local shape features (neural network texturization; NNT). Image information (Fig. 3d) was strongly reduced by both texturizations (Original vs WT: p = 6*10^−8^; Original vs NNT: p = 0.0007), but more so by wavelet texturization (WT vs NNT, p = 8*10^−5^). Cross-information analysis revealed that neural representations of texturized images significantly overlapped with those of original stimuli (Fig. 3d dark gray bar; cross-information > 0; WT: p = 0.012; NNT p = 0.0009). However, this overlap was small compared to the overlap expected for identical representation (Fig. 3d light vs. dark gray bar; cross-information < expected; WT: p = 8*10^−6^; NNT: p = 2*10^−7^) indicating that neural representations generalized only weakly.

Category information (Fig. 3e) was more drastically reduced by texturization than image information and the neural responses to categories were less discriminable for the WT stimuli retaining no shape features than for the NNT stimuli retaining local shape features (WT vs NNT p = 3*10^−5^). Nonetheless, there was still significant category information for both kinds of texturized stimuli (NNT > 0: p = 4*10^−7^; WT > 0: p = 0.013) and these category representations were to some extent shared with those evoked by original stimuli (Fig. 3e dark gray bar; cross-decoding > 0: NNT: p = 9*10^−6^; WT: p = 0.0001). Again, this generalization was only partial, as it was smaller than expected for identical representations (Fig. 3e light vs. dark gray bar; cross-information < expected; NNT: p = 1*10^−6^; WT: p = 0.053).

Finally, we directly confirmed that both texturizations strongly changed neural responses to individual images (Fig. 3f, left, WT: p = 8*10^−8^; NNT: p = 4*10^−8^). These representational shifts were significantly collinear both within (Fig. 3f, middle, WT: p = 1*10^−7^; NNT: p = 6*10^−8^) and across categories (Fig. 3f, right, WT: p = 2*10^−7^; NNT: p = 3*10^−7^). This indicates that shape features were to some extent shared by natural images both within the same, but also across categories.

In sum, we found that both image and category information were reduced by texturization. In both cases, more information was retained for neural-network texturized stimuli due to the preservation of local shape features. The strong, but not complete reduction of category information indicates that shape features were a prominent, albeit not the only driver of category information.

### Shape and spectral power predominantly account for category selectivity

Neither differences in spectral power nor in global or local shape alone could account for neural image and category information. Thus, we next assessed the joint effect of both features on the representation of natural images, by comparing neural responses to power-matched stimuli (Original PM) with those of texturized and power-matched stimuli (NNT PM and WT PM) (Fig. 3g, h, i). Importantly, any imageor category selectivity preserved after texturization and power-matching would have to be due to the only remaining features of the stimuli, which are the local spatial correlation structure (for both NNT PM and WT PM), local shape (only in the case of NNT PM), or retinotopy (in the case of image information for both NNT PM and WT PM).

Removing shape features by texturization and power matching further reduced both image (Fig. 3g, Original PM vs WT PM: p = 2*10^−5^, Original PM vs NNT PM: p = 0.002) and category information (Fig. 3h, Original PM vs WT PM: p = 6*10^−8^, Original PM vs NNT PM: p = 2*10^−7^). Image information about spectrally matched original and texturized images peaked at the same magnitude but differentially developed over time. We found no significant cross-information about images (Fig. 3g, dark gray bar; cross-information > 0; WT PM: p = 0.83; NNT PM: p = 0.16). Category information (Fig. 3h) was almost entirely removed by the removal of both shape and spectral power features, but remained significant (Fig. 3h, NNT PM > 0: p = 0.0006; WT PM > 0: p = 0.047). Furthermore, there was a significant generalization of category information between the neural responses to original and global-shape-removed, spectrally matched stimuli (Fig. 3h, dark gray bar; cross-information Original PM vs. NNT PM > 0; p = 1*10^−5^). This cross-information did not deviate from the expectation for identical representations (Fig. 3h, dark vs light gray bar; cross-information < expected; p = 0.24), indicating that the residual category representations were largely shared. In contrast, we found no generalization between the neural responses to original and fully shape-removed, spectrally matched stimuli (Fig. 3h, dark gray bar; cross-information Original PM vs. WT PM > 0, p = 0.83).

Like our previous findings, both texturization algorithms also strongly changed the neural responses to individual images when they were spectrally matched (Fig. 3i, left; WT PM: p = 8*10^−8^; NNT PM: p = 1*10^−7^). Again, these representational shifts were significantly collinear both within (Fig. 3i, middle; WT: p = 3*10^−7^; NNT: p = 4*10^−8^) and across categories (Fig. 3i, right; WT: p = 2*10^−6^; NNT: p = 1*10^−6^). This suggests that shared shape features of natural images persisted after the removal of global spatial frequency and orientation energy.

In sum, these findings show that spectral power and shape features together nearly completely accounted for the neural selectivity for categories of natural images. While the remaining local shape features in global-shape-removed stimuli elicit some generalizable category information, the textural correlation structure preserved after the removal of all shape only carried marginal category information.

### Retinotopic shifts contribute to image information

All modifications discussed so far involved changes in retinotopic patterns – changes in the spatial layout of light intensities and local features. Likewise, different natural images within the same modification also differ in their retinotopic patterns. To isolate the effects of retinotopy, we applied the two texturization algorithms twice to the same power-matched images, using different random seeds (Fig. 1c; WT 1 PM, WT 2 PM, NNT 1 PM, NNT 2 PM). The seed manipulation selectively changed retinotopic patterns, because the images did not contain shape features, while spectral power as well as texture was the same for both seeds.

First, we confirmed that repeated, independent texturizations lead to image information of similar magnitude (Fig. 3k, WT 1 PM vs WT 2 PM: p = 0.13, NNT 1 PM vs NNT 2 PM: p = 0.49). However, image representations generalized only to a small extent (Fig. 3k, dark gray bars; cross-information > 0, WT: p = 0.04; NNT: p = 0.0001) and significantly less than expected for identical representations (Fig. 3k, dark vs light gray bars; cross-information < expected; WT: p = 3*10^−5^; NNT: p = 1*10^−5^). Thus, the neural representations of repeated texturizations of the same image were largely distinct despite the preserved local shape and/or textural correlation structure.

Category information was low for both random seeds, but significantly different between the two seeds for neural network texturization (Fig. 3m, WT 1 PM vs WT 2 PM: p = 0.55; NNT 1 PM vs NNT 2 PM: p = 0.018). Interestingly, category information significantly generalized for neural network texturization but not for wavelet texturization (Fig. 3m, dark gray bars; cross-information > 0, WT: p = 0.62; NNT: p = 3*10^−7^), and this generalization was not smaller than expected for identical representations (Fig. 3m, dark vs light gray bars; cross-information < expected; WT: p = 0.24; NNT p = 0.99).

Consistent with the small generalization of image information between texturizations, neural responses to individual images were significantly different between random seeds (Fig. 3n, left; WT: p = 2*10^−5^; NNT: p = 8*10^−6^). As the two seeds were independent, they led to random shifts in representational space, which were neither collinear within nor across categories (Fig. 3n, middle and right; WT: within p = 0.14; across p = 0.91, NNT: within p = 0.89; across p = 0.63).

In sum, independent texturizations of the same images resulted in similar amounts of both image and category information. In contrast to image information, the remaining category information about neural network texturized images generalized across random seeds, indicating that it was driven by local shapes or textural correlation structure.

### Decomposing the contribution of stimulus features to neural information

The employed stimulus space and the above analyses afforded the possibility to decompose neural information about natural images and semantic categories into their constituent parts explained by five different visual features and to quantify the dynamics of their unique contributions: retinotopy, global shape, local shape, spectral power, and textural correlation structure (Fig. 4). Retinotopic information can be defined as the neural separability of two images that differ only in the spatial distribution but not in the marginal statistics of their features. We thus quantified retinotopic information as the average representational distance between two independent wavelet-texturizations of the same image (Fig. 3n left; overall). Our stimulus space only allowed to compute the contribution of retinotopy for image information. We chose not to include further modifications allowing for an estimation of category retinotopy due to the assumption that retinotopy could only be marginally informative about category membership, which should mostly abstract over retinal position.

**Fig 4.**
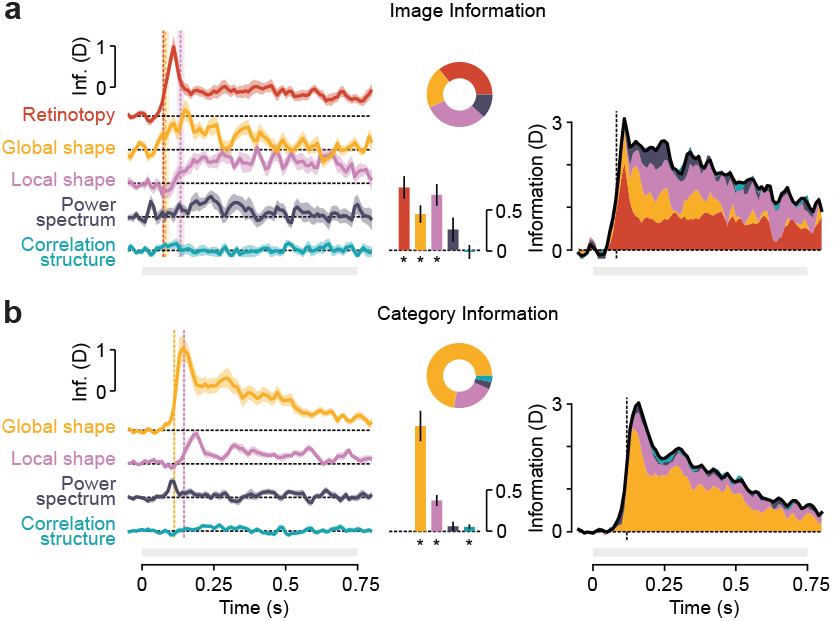
Contribution of stimulus features to neural information. (**a**) Left: Time course of 5 components of image information. Vertical dashed lines indicate the latency of half maximum information. Middle: The colored bars show the average component information 0.15 to 0.5 s post stimulus onset. Asterisks below bars indicate significant information (p < 0.05). The pie chart indicates the components’ relative contributions to image information. Right: Stacked timecourses of the 5 components of image information. The vertical dashed line indicates the latency of half maximum total image information. (**b**) as in (a) for 4 components of category information. All error bars indicate the SEM across subjects.

Global shape information is the neural information lost by removing the global cohesion of local patches of shape, which we computed as the difference between neural information of original and neural-network texturized stimuli. We define local shape information as the neural information lost by removing local shape patches from the stimuli, that is, the difference between the neural information about neural-network and wavelet texturized stimuli. Spectral power information was defined as the neural information lost by power-matching stimuli, and here computed as the difference in neural information about wavelet-texturized and wavelet-texturized power-matched stimuli. We chose to use wavelet-texturized stimuli for the evaluation of this parameter to avoid possible interactions with the global and local shape information in original or neural-network texturized stimuli.

Finally, the contribution of textural correlation structure can be quantified by assessing the neural information about wavelet-texturized images, after controlling for the effect of retinotopy. In the case of image information, we thus computed the contribution of the correlation structure as the difference between the neural information about wavelet-texturized stimuli and retinotopic information as defined above. In the case of category information, the correlation structure contribution was defined as the neural information about wavelet-texturized stimuli, as any category-specific retinotopic information would have been removed by the manipulation. Notably, these five components, by definition, add up to the total image and category information. Using this decomposition, we found a significant retinotopic contribution to image information (Fig. 4a, p = 2*10^−5^ 0.15 to 0.5 s) that showed a steep rise (half-maximum latency: 75 ± 2 ms). This component was largely responsible for the characteristic onset shape of image information and subsequently decreased to a constant level. In contrast, significant global (p = 0.0003) and local (p = 4*10^−5^) shape contributions to image information rose more slowly. Global shape information had a similar latency as retinotopy information (half-maximum latency global shape 82 ±13 ms; difference to retinotopy p = 0.6), while local shape information was significantly later than retinotopy and global shape information (half-maximum latency local shape 135 ± 14 ms; difference to global shape and retinotopy both p < 0.001) (Fig. 4a). The contribution of spectral power to image information was only marginally significant (p = 0.051) and the contribution of correlation structure to neural image information did not reach significance (p = 0.59).

For neural information about stimulus category, global shape was by far the largest significant contributor (p = 1*10^−6^) with a strong onset response and slow decline (Fig. 4b). Analogous to image information, local shape contributed significantly (p = 2*10^−5^) but showed a significantly delayed time course compared to global shape (half-maximum latencies: global shape 113 ± 4 ms; local shape 147 ± 5 ms; difference: p = 8*10^−6^) (Fig. 4b). The contribution of spectral power was not significant (p = 0.15) and the contribution of correlation structure was small (p = 0.047).

### Category specificity of neural information

Our stimulus space contained images from a wide range of categories. Thus, we next quantified image and category information as well as the effect of image modifications separately for different categories.

For original stimuli, we observed striking differences in both image and category information across categories (Fig. 5a, b, left). While pictures of animals and artificial objects evoked the most image information, human faces led to the lowest image information (Fig. 5a, left). Conversely, human faces evoked the highest category information (Fig. 5b, left). While these results may be partially explained by differences in stimulus variability (e.g., different artificial objects vs. frontally depicted human faces), they may also reflect a specialization of the human visual system. Removing overall spectral power (Original PM, Fig. 5a, b, middle) removed some image and category information, while further removing global and local shape (WT PM, Fig. 5a, b, right) removed all category information while retaining image information that was not category specific.

**Fig 5.**
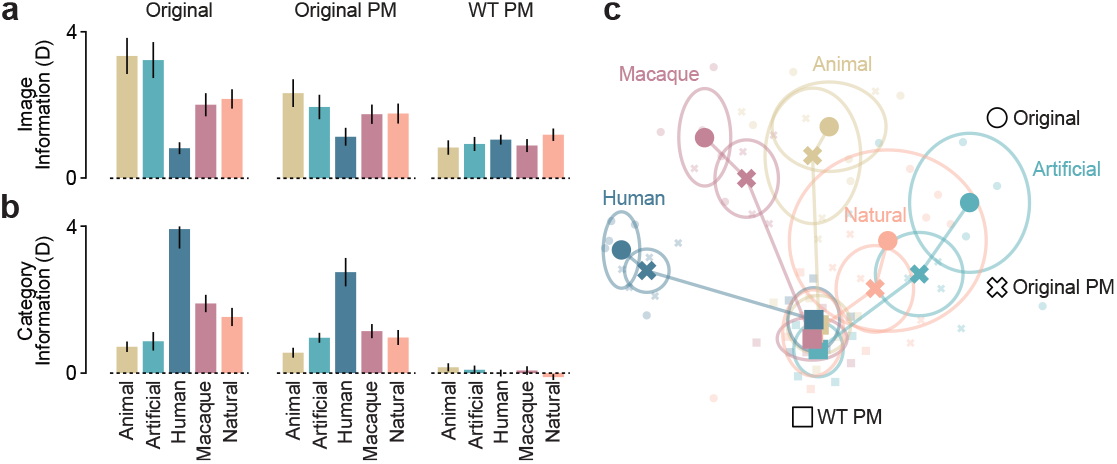
Neural information by category and modification. (**a**) Image information by category for original, power matched and wavelet texturized power matched stimuli 0.15 to 0.5 s post stimulus onset. All error bars indicate the SEM across subjects. (**b**) as (a) for category information. (**c**) Multidimensional scaling (MDS) of the neural representational space into a 2dimensional space for original, power matched and wavelet texturized power matched stimuli. Small transparent markers indicate all individual images. Large opaque markers indicate the average across images per category and modification. The circles indicate the 2-dimensional standard deviation of the position of images per category and modification.

An ANOVA statistically confirmed these results. We found categorical differences in image information for all, but wavelet-texturized power-matched stimuli (originals: p = 1*10^−10^; original PM: p = 0.0006; NNT PM: p = 0.02; WT PM: p = 0.45). For category information only original and power-matched originals showed significant differences between categories (originals: p = 3*10^−15^; original PM: p = 3*10^−13^; NNT PM: p = 0.19; WT PM: p = 0.48). Overall, this showed that category selection strongly influenced image and category information and illustrates how ordinate specificities disappear with the removal of object shape and spectral power differences.

### Representational geometry of stimulus modifications

All above results describe differences between the neural activity elicited by different images, semantic categories, and changes in this high-dimensional representational space due to modifications of these stimuli. To provide a summary, and intuitive interpretation of the complex effects described so far, we created a two-dimensional embedding of the full representational space using multidimensional scaling (MDS) (Fig. 5c).

We selected original, power-matched and wavelet-texturized power-matched stimuli to illustrate the effect of spectral matching and texturization on image and category representations. The neural activity patterns elicited by original images within each category were distinct, as evident by their wide distribution. Notably, neural responses to human faces were farthest away from the other categories and showed the smallest within-category variability, in line with the above finding of highest category information for human faces (Fig. 5b, left). Matching power spectra between images (Original PM) moved neural responses to different stimuli closer together, both within and across categories. Finally, combined wavelet texturization and spectral power matching (WT PM) further decreased variability within categories, and abolished variability across categories. In sum, this visualization shows how the progressive removal of relevant image features collapses the representational space, and neural activity patterns converge onto a single point, thereby greatly reducing image information and removing category information.

### Invariance across stimulus sets

In a final set of analyses, we tested if the selection of specific images for each set of 25 original stimuli had a fundamental effect on our results. To this end, we compared the two groups of subjects that viewed distinct sets of stimuli. We pooled original and power-matched stimuli, and performed an attenuation-corrected representational similarity analysis [25,26] of neural information 0.15 s to 0.5 s post stimulus onset for all stimuli of different categories between stimulus sets. We derived subject-specific estimates using jackknife resampling. We found no significant difference (two-sided t-test, p = 0.58) between the reliability of neural information within set 1 (0.95±0.03) and within set 2 (0.92±0.05). Furthermore, the attenuation-corrected correlation coefficient, which indicates the similarity of category differences across stimulus sets after accounting for the reliability within each set, was not significantly lower than 1 (1.02±0.01, p = 0.7). Thus, there was no difference in data quality, and the representational space spanned by the five categories generalized across the two stimulus sets.

## Discussion

The selectivity of neural responses for natural visual stimuli and semantic categories has been an area of intense research [1,2,27–34]. Selective responses have been identified across many brain areas and recording modalities, most prominently in object-selective visual and temporal cortices. Nonetheless, owing to the complexity of natural images, it remains unclear to what extent different visual features underly neural selectivity and support human object perception. Here, we address this gap by characterizing the contributions of retinotopy, textural correlation structure, spectral power, and local as well as global shape to image- and category selectivity in human MEG data. Using a stimulus space consisting of natural images from five categories, as well as several modified versions of each image, we found that all modifications disrupting specific visual features reduced neural information about both images and categories. By assessing the changes in representational geometry and information content induced by each modification, we could quantify the contributions of all five features to overall selectivity and determined that retinotopy was the strongest driver of image information, while global shape most prominently influenced category information. These findings contribute to our understanding of natural stimulus processing in the human brain, and provide a basis for understanding the sources of image- and category selectivity reported across the literature [1,12,30]. More generally, our results can help constrain computational models of human object recognition and illustrate how a stimulus space covering the range of visual features can be a valuable tool to decompose neural responses. We anticipate that similar manipulations will be crucial to scrutinize the behaviour of human observers as well as neural network models.

### Dissociating sensory and semantic discriminability

Previous research has shown neural selectivity for many sensory features as well as semantic categories of natural stimuli [14,16–20,35–37]. An implicit assumption of studies separating the neural discriminability of images within a category from that of categories is that the former is driven by sensory features, whereas the latter reflects semantic meaning. In its extreme form, this assumption is untenable: categorization depends on reliable perceptual differences [38], and semantic meaning varies within each possible category. This makes it challenging to cleanly differentiate sensory from semantic contributions, and even sophisticated experimental designs rigorously controlling for the effect of specific features rely on perceptual differences for observers to successfully categorize stimuli [17,18].

Here, we chose a different approach: rather than attempting to dissociate sensory and semantic aspects of neural image representations, we quantify the contribution of all constituent visual features; regardless of whether these contributions are sensory or semantic in nature. Thus, the neural responses driven by each feature may subsume both purely bottom-up processing, and thereby the contribution to stimulus or category selectivity due to different feature values, and any subjective processing elicited by or associated with the feature. Importantly, we still assess selectivity for both individual images and stimulus categories. However, we quantify the contribution of multiple sensory features to both types of selectivity.

### Decomposing feature contributions to neural information

Decomposing neural information into its constituent parts driven by different visual features required several methodological choices. First, we assessed the contribution of retinotopy using two sets of stimuli which were power-matched and texturized, and thus, did not meaningfully vary in global or local shape and spectral power. This allowed us to estimate changes in neural responses due to retinotopy. However, it prevented the estimation of any contribution of retinotopy to neural category selectivity. Here, we made the simplifying assumption that retinotopy may be a strong driver of image information individual images strongly differ in where their elements are - but much less of category information, as the nature of categories as grouping multiple members and generating invariance over the exact spatial location should largely imply an abstraction over retinotopy effects. Nonetheless, retinotopy may theoretically contribute to category information - where a certain pattern of light is detected on the retina may be informative about the semantic category of its source [39,40]. Thus, further studies may be able to quantify contributions of retinotopy to category selectivity. In our results, these contributions may be included in the variability explained by some of the other features, specifically global and local shape. Furthermore, our approach rests on the assumption that retinotopic differences are approximately of equal magnitude in natural and texturized stimuli.

Second, we partially decompose neural information by subtraction. The neural information about natural stimuli within a certain modification and category is related to the variability of the underlying neural patterns. As variances are only additive for independent variables, an accurate decomposition relies on each feature contribution having an independent effect on the neural representation of natural stimuli. In other words, the effects of different features should not be collinear. Indeed, it is unlikely that two modifications have perfectly collinear effects on the neural representation of a stimulus: the modified features themselves are not redundant and cannot be expected to be perfectly correlated – knowing the global shape of a natural stimulus is not sufficient to predict its local shape or spectral power spectrum. Furthermore, sensory neural representations of different features are at least partially orthogonal and driven by distinct populations. On the other hand, some collinearity is plausible, especially when neural representations reflect the semantics of presented stimuli: any modification degrading a natural stimulus may weaken the neural representation of its meaning, and thus lead to a representational shift towards a common origin for all stimuli. In sum, it is reasonable to assume that the effects of modifications in our stimulus space are largely independent, but partially collinear. This partial collinearity could lead to small deviations of estimated feature contributions. Future studies using similar stimulus spaces allowing to compute feature contributions in different sequential orders, or to directly assess the effect of retinotopy on unmodified stimuli may mitigate this.

Importantly, and despite these limitations, our approach allows to comprehensively attribute components of neural information to specific stimulus features in such a way that these components together fully explain the neural information at hand.

### Relative contributions to image and category information

Unsurprisingly, we found that each feature influences neural image selectivity. Applying any modification to our set of natural stimuli subjectively led to clear perceptual differences, which should accordingly be reflected in neural activity. Whether any set of degraded natural stimuli is expected to support neural selectivity for categories is less clear. We found that both global and local shape had sizeable effects on category selectivity, as evidenced by the loss in category information due to texturizing stimuli; with global shape explaining the bulk of category information. This is consistent with previous findings showing a strong contribution of global-shape-like features in fMRI [18,41], M/EEG [17,19,42] or electrophysiological recordings from area IT in non-human primates [43]. In our data, global shape information temporally preceded local shape information. This accords well with the idea of an initial processing sweep extracting the gist of a natural image [39,40]. It is, however, important to note that the concept of gist has previously been applied to natural scenes rather than images of single objects. Furthermore, while gist captures global summaries of image features, it does not perfectly correspond to any of the fundamental features defined in the present study and rather constitutes an alternative parametrization of the visual feature space.

Interestingly, it has been suggested that object-selective areas in the ventral visual stream do not explicitly represent object-centered, viewpoint-invariant global shape [44], and that local visual features are the main drivers of object- and category selectivity. This is in contrast with our finding of global shape as the most prominent contributor to neural category information.

How can these findings be reconciled? Notably, in our results, the combined effect of retinotopy, local shape, and spectral power on image information is still larger than the effect of global shape alone, consistent with a dominance of broadly defined texture selectivity over global shape selectivity [45]. Secondly, in the present study we did not differentiate between abstract, view-invariant shape and global visual shape features such as contour or curvature at a large spatial scale. Thus, these latter visual features may contribute to shape-associated category selectivity both in previous studies and in our results [44,46,47].

In a similar vein, it has been found that while neural responses in human object-selective cortex and monkey IT support the categorization of natural stimuli, they do not allow differentiating well between natural and texturized stimuli [48]. This has been interpreted as evidence that these ventral stream areas encode textural features allowing for flexible object categorization rather than explicitly representing objects. Interestingly, we here find that the neural representations of natural and texturized images are well separated, as evidenced by the significant collinearity of representational shifts both within and across categories. This suggests that either these object-selective areas are more strongly selective for a natural feature arrangement than previously shown, or selectivity in our results was driven by other areas.

While shape features were the main drivers of category selectivity in our data, they did not fully explain category information. Previous research has shown that there is shape-independent category information [17,18,37], which was interpreted as genuine, semantic category information. Here, we find that neither neural-network texturization nor, more importantly, wavelet-based texturization fully abolished category information, suggesting that the neural representation of the local spatial correlation structure is still slightly selective for semantic categories. In line with this, we found a distinct representation of correlation structure when assessing the cross-information about natural images across repeated texturizations. Notably, this effect was not identified in either the decomposition analysis or the cross-information about image categories, which is likely due to the small effect size and the partially independent nature of these tests. For this reason, only the cross-information test (using data from both seeds, rather than just one, and image information which was generally higher than category information) may have been sensitive enough. In sum, our results support the careful conclusion that correlation structure contributes to the selectivity for natural images and may carry some semantic information.

The effect of spectral power on both image and category information was small and did not reach significance in our main analysis which computed the spectral power contribution after accounting for all other features. A less conservative approach directly assessing the effect of power-matching on otherwise unmodified stimuli may reveal a larger contribution of spectral power. However, this could be driven by partial dependencies between the overall spatial frequency and orientation energy described by the power spectrum and the other visual features. For example, global and local shape may become less recognizable when powerspectra are matched across stimuli.

Overall, both image and category information were largely driven by spatially located shape information – global and local shape as well as retinotopy, and much less so by marginal summary statistics such as the overall spectral energy and correlation structure. This is in line with the fact that even clear visual features presented in isolation are challenging to decode [11,49,50]. This may reflect the retinotopic organization of visual cortex, implying that where something is, is easier to read out than what it is.

### Tradeoff between exemplar and category discrimination

Object recognition serves both the rapid identification of category membership, such as telling a tiger from a tree, and the fine discrimination of individual category members, such as different human faces. These tasks require a conflicting optimization of the neural representational space: category discrimination benefits from closely grouping category members together while maximizing the distance between categories, whereas exemplar discrimination requires variable representations within a category.

This tradeoff is also apparent in our data: First, image and category information are anticorrelated across categories, such that categories with closely grouped exemplars are more distinguishable from other categories. Most notably, the neural representations of human faces are similar, but far from all other categories. This may be due to the exceptional significance of conspecifics, which can lead to a gradient of response magnitudes according to semantic similarity to humans [51,52]. When visual features are degraded, differences between categories are diminished. Ultimately, simultaneous power-matching and wavelet texturization largely abolishes differences in the amount of image information between categories, and at the same time also category information itself. This implies that the magnitude of the remaining within-category differences in retinotopy and local correlation structure is not category specific.

### Summary and conclusion

In conclusion, we decomposed the stimulus and category selectivity of neural responses to natural images in the human brain, and quantify the contributions of retinotopy, global and local shape, the spatial power spectrum, and the textural correlation structure. Our results show how these features jointly account for neural selectivity. This comprehensive assessment provides the basis for a better understanding of the neural mechanisms underlying human object recognition.

## Acknowledgements

This study was supported by the European Research Council (ERC; https://erc.europa.eu/) CoG 864491 (M.S) and by the German Research Foundation (DFG; https://www.dfg.de/) project 276693517 (SFB 1233) (M.S.).

## Author contributions

JKS: Conceptualization, Methodology, Data curation, Formal analysis, Visualization, Writing – original draft, Writing – Review & Editing; MSh: Investigation, Data curation, Formal analysis, Writing – Review & Editing; FS: Conceptualization, Supervision, Methodology, Writing – original draft, Writing – Review & Editing; MSi: Conceptualization, Supervision, Resources, Project administration, Funding acquisition, Visualization, Writing – Review & Editing

## Materials and Methods

### Participants

18 healthy volunteers (10 male, mean age 27) with normal vision participated in this study and received monetary reward. All subjects except for one were right-handed. Participants provided written informed consent prior to the start of the experiment. The study was conducted in accordance with the Declaration of Helsinki and was approved by the ethical committee of the Medical Faculty and University Hospital of the University of Tübingen.

### Behavioral task & stimuli

Participants performed a visual attention paradigm with passive fixation. On each trial, they had to maintain central fixation while a stream of stimuli was presented (edge length: 10 degrees of visual angle, 0.75 s initial blank fixation baseline followed by 0.75 s per image). With the final image of the sequence, a visual attention target appeared, instructing the subjects to lift their dominant hand’s index finger. If no response was given within 0.75 s, the target was counted as missed. Fixation was controlled continuously within a circular window with 2 degrees radius. Fixation breaks and eye blinks led to the trial ending prematurely.

Between 2 and 20, with a mean of 12, images made up the stimulus stream in each trial. The stimulus count was sampled from a Poisson random distribution, creating a uniform hazard rate; and cue location was selected randomly from 40 equidistant locations distributed across 4 radially equally spaced eccentricities covering the displayed stimulus. Thus, the appearance of the target was unpredictable in space and time.

The task was run using Psychtoolbox-3 [53,54] and custom code in MATLAB 2017b.

### Setup & recording

We recorded MEG (Omega 2000, CTF Systems, Inc., Port Coquitlam, Canada) with 275 channels at a sampling rate of 2,343.75 Hz in a magnetically shielded chamber. Participants sat upright in a dark room, while stimuli were projected onto a screen at a viewing distance of 55 cm using an LCD projector at 60 Hz refresh rate. We recorded eye movements using an Eye-link 1000 system (SR Research, Ottawa, Ontario, Canada). Two sessions were recorded per subject. One session of one participant had to be discarded for technical reasons.

### Stimulus space and image modifications

For each participant, we used a stimulus space consisting of 200 grayscale images. 25 natural stimuli were selected from 5 different categories (animal, artificial, human, macaque and natural) with 5 exemplars each. Animal images were full-body shots of different mammals on a naturalistic background selected from the THINGS database [13]. Artificial objects were images of varying man–made objects taken, also selected from the THINGS database. Human images were realistic, artificially generated male and female portrait shots, created using a generative adversarial model [55], https://thispersondoesnotexist.com/). Macaque stimuli were comprised of body photographs, with the face turned towards the camera. Natural images were of various fruit and vegetable items, taken from the THINGS database. The original images were modified using two available open-source texturization algorithms, one based on wavelet decomposition [22] and a convolutional neural network based approach [21]. Wavelet texturization was performed with default parameters [56]. Spectral differences between images and categories were masked by matching the 2D Fourier power spectrum of each image to a common mean (power match, PM) computed from images of the THINGS database. Texturizations were performed twice to generate different version of the image with differing spatial positioning of image features. These repetitions were only shown in their power-matched version. In a final step, all images had their pixel luminance distributions matched to a common mean distribution and were gamma corrected. This created a stimulus space with the following 8 modifications: Original, Original PM, Neural Network Texture 1, Neural Network Texture 1 PM, Neural Network Texture 2 PM, Wavelet Texture 1, Wavelet Texture 1 PM, Wavelet Texture 2 PM. This stimulus space was generated twice, with different original images, creating 2 sets of matching stimuli thar were randomly assigned to subjects, to control for confounds due to stimulus selection.

### Preprocessing

Data was preprocessed using MATLAB and the Fieldtrip toolbox [57]. Infrequent SQUID channel jumps occurred in one session and were interpolated using 150 samples on either side of the jump. Data were lowpass filtered at 30 Hz (Butterworth, 6th order, zero-phase), resampled to 100 Hz, and epoched into sub-trials with respect to each stimulus onset. Each subtrial was baseline-corrected, using the preceding 250 ms time window. The initial and final stimuli of each trial were excluded to avoid confounds due to the initial onset of the stimulus stream or the attention target. Stimulus presentations with an average MEG signal larger than 8 standard deviations from the mean across all sub-trials were removed from the analysis.

### Multivariate pattern analysis

We computed pairwise neural dissimilarities between the neural responses to all stimuli using Cross-validated Mahalanobis distances (CVMD, [23], a two-fold cross-validation scheme and spatial pre-whitening using shrunk covariance estimates [58] of the neural responses to all included stimuli. Cross-validation was repeated 200 times with different random seeds to decrease dependencies on data partitioning. Dissimilarity matrices were averaged over these repetitions.

### Category information

Category information can be extracted by pairwise training and testing on different categories, using non-identical image pairs. As this is computationally demanding, we employed a mathematical shortcut leading to the same results and extracted category information from the neural dissimilarity matrices. To do so, we computed the mean pairwise across categories as well as the mean pairwise distance within categories. Category information was then defined as the across-category distance minus the within-category distance.

### Cross-information between modifications

We estimated the extent of neural representational overlap between modifications using cross-information, by training on neural responses to stimuli from one modification and testing on stimuli from another. As the maximally possible amount of shared information between modifications depends on the information available about stimuli in each individual modification, we compared the strength of the shared across-modification representation to the strength of both modifications’ representations. [24,59]. To this end, we computed the expected cross-information

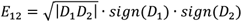

where D_1_ and D_2_ denote CVMD between an image pair in the two modifications. If stimulus representations were identical in both modifications, differing only in magnitude, the cross-information D_12_ would approach E_12_, which constitutes a lower bound of the true expected cross-information between modifications.

### Collinearity computation

We assessed the effect of stimulus modifications on individual images, members of the same category, or all images. To do so, we first computed the dissimilarity between any stimulus and its modified version, averaged over all 25 stimuli, which we refer to as the overall representational shift. Then, we computed the collinearity between the representational shifts due to modifying all stimuli within the same category. Here, we used the same mathematical equality as we did for deriving category information and defined the within-category collinearity as the mean dissimilarity between stimuli of the same category but different modifications minus the mean dissimilarity of stimuli of the same category and the same modification. This mathematically corresponds to the cross-information about the respective modification across stimuli of the same category. Similarly, we defined the across-category collinearity as the mean dissimilarity between stimuli of different categories and different modifications minus the mean dissimilarity of stimuli of different categories but the same modification. This mathematically corresponds to the cross-information about the respective modification across stimuli of different categories.

### Feature contribution decomposition

To quantify the contributions of different visual features to the neural responses to natural stimuli, we decomposed overall image and category information. First, we computed retinotopic information as the average representational shift between the two versions of each wavelet-texturized, power-matched stimulus:

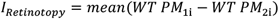

Retinotopic Information exists only for image information, as category information generalizes across retinotopic changes. For further computations, the retinotopic information of category information can be thought of as 0.

Then, we computed the information about global shape (I_G_), local shape (I_L_), the power spectrum (I_P_) and the correlation structure (I_C_) as the difference between the average neural information in two modifications. These modification contrasts were chosen such that stimuli differed in the presence of the respective feature of interest only:

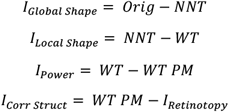

This procedure was identical for image and category information, apart from the computation of I_Retinotopy_, which was omitted for category information, and I_Corr Struct_, which was equal to WT PM, that is, the category information about wavelet-texturized, power-matched stimuli, due to the absence of a retinotopy contribution.

### Multidimensional Scaling

Multidimensional scaling of the pairwise decoding values into a 2-dimensional space was done on data from Original, Original PM and WT PM stimuli averaged across a time window from 0.15 s to 0.5 s The scaling was computed using the MATLAB function mdscale.

### Latencies

We estimated the latency of information as the time at which information reached half its maximum from 0 to 0.25 s post stimulus onset. We estimated the latencies of feature contributions to information for those contributions that were significant and showed a discernable onset dynamic (image information: retinotopy, global shape, local shape; category information: global shape, local shape). For all latency estimates, information time-courses were smoothed with a Hanning window (100 ms full width at half maximum). For statistical comparisons of latencies between types of information we estimated the standard error of latencies by bootstrap across subjects (1000 resamples). We then assessed the significance of latency differences using t-statistics.

### Statistics

All statistics were computed across subjects. Statistical tests were performed against an alpha-value of 0.05 for the average information from 0.15 to 0.5 s post stimulus onset. All bar plots display the same interval. All error bars indicate the standard error of the mean across subjects. Statistical significance between cross-decoding and expectation values and for information larger than zero was determined using one-sided t-statistics. All other significances were assessed with two-sided t-statistics.

### Software

Preprocessing and data analysis was performed using custom code using Matlab2017b and the Fieldtrip toolbox [57], as well as Python 3.9, with dependencies in scikit-learn 1.1.2 [60], NumPy 1.22.3 [61], Pandas 1.4.4 [62], Pingouin 0.5.2 [63].

